# Scene wheels: Measuring perception and memory of real-world scenes with a continuous stimulus space

**DOI:** 10.1101/2020.10.09.333708

**Authors:** Gaeun Son, Dirk B. Walther, Michael L. Mack

**Author notes:** Correspondence concerning this article should be addressed to Michael L. Mack, Department of Psychology, University of Toronto, Sidney Smith Hall, 100 St George St, Toronto, ON M5S 3G3, Canada.

## Abstract

Precisely characterizing mental representations of visual experiences requires careful control of experimental stimuli. Recent work leveraging such stimulus control has led to important insights; however, these findings are constrained to simple visual properties like colour and line orientation. There remains a critical methodological barrier to characterizing perceptual and mnemonic representations of realistic visual experiences. Here, we introduce a novel method to systematically control visual properties of natural scene stimuli. Using generative adversarial networks (GAN), a state-of-art deep learning technique for creating highly realistic synthetic images, we generated scene wheels in which continuously changing visual properties smoothly transition between meaningful realistic scenes. To validate the efficacy of scene wheels, we conducted two behavioral experiments that assess perceptual and mnemonic representations attained from the scene wheels. In the perceptual validation experiment, we tested whether the continuous transition of scene images along the wheel is reflected in human perceptual similarity judgment. The perceived similarity of the scene images correspondingly decreased as distances between the images increase on the wheel. In the memory experiment, participants reconstructed to-be-remembered scenes from the scene wheels. Reconstruction errors for these scenes resemble error distributions observed in prior studies using simple stimulus properties. Importantly, perceptual similarity judgment and memory precision varied systematically with scene wheel radius. These findings suggest our novel approach offers a window into the mental representations of naturalistic visual experiences.

---

The ultimate goal for visual cognition research is understanding how people encode the complexities of everyday visual experience into memory traces that can be retrieved moments to hours to days later. A guiding principle of this research is the use of simple stimuli (e.g., spatial frequency, orientation, or colour) to investigate various aspects of visual representation including the capacity of iconic memory (Bradley & Pearson, 2012; Sperling, 1960), the storage structure of visual working memory (VWM; Bays et al., 2009; Brady & Alvarez, 2011; Luck & Vogel, 1997; Zhang & Luck, 2008), and decay or bias of visual long-term memory (Magnussen et al., 2003; Ester et al., 2020). Although decades of this research have characterized many facets of the visual memory and perception system, it remains an important open question how these findings based on simple visual stimuli generalize to memory for realistic visual environments. The key limitation preventing such advances is the methodological hurdle of proper empirical control of realistic scene stimuli.

The careful control of the physical properties of stimuli is a critical element for many domains of cognitive psychology. By systematically manipulating physical properties of stimuli, corresponding changes in task performance can be identified and leveraged to characterize changes in cognitive ability. One prominent example is the change detection paradigm in VWM studies in which participants have to detect any differences between two visual arrays presented sequentially. By manipulating the physical properties of stimuli composing the visual arrays, researchers can identify those features that are successfully encoded and remembered (Harrison & Tong, 2009; Kiyonaga & Egner, 2016; Johnson et al., 2009; Lin & Luck, 2009; Luck & Vogel, 1997; Wilken & Ma, 2004). For example, Lin & Luck (2009) manipulated the similarity of target and foil object colours by measuring coordinate distance between the stimuli in CIE (Commission Internationale de l’Elcairage) colour space (Smith & Guild, 1931). Based on this deliberate manipulation, they demonstrated that visual working memory (VWM) performance increases when target and foil colours are similar, a finding that speaks to a fundamental relationship between a pervasive visual feature and how the visual memory system encodes it.

The level of stimulus control available for a feature like colour is, however, very difficult to achieve in the real-world scene domain. Whereas colour spaces like CIE offer well-established links between physical and perceptual similarity, the amount of information and complexity in realistic scenes makes it difficult to define the similarity between any two scenes in a manner that reflects human perception. One recent study (Konkle et al., 2010) took an indirect approach to overcoming this challenge by manipulating the semantic similarity of target and foil scenes as a proxy for visual similarity. The underlying assumption was that scene exemplars would share similar visual properties to some extent if they are in the same semantic category (e.g., two kitchens are more visually similar than a kitchen and bedroom). Certainly, many natural categories exhibit family resemblance with shared visual properties; however, such semantic manipulations are a noticeable departure from the deliberate manipulation of visual properties like colour or orientation (Harrison & Tong, 2009).

The fine control of stimulus properties is essential for adapting new experimental paradigms. Continuous report is a recently developed paradigm that makes it possible to more directly observe a representational structure of the visual system (Bays & Husain, 2008; Wilken & Ma, 2004; Zhang & Luck, 2008). In this paradigm, participants select the stimulus they experienced using a highly controlled response scale that contains all possible stimulus options used in the study and is organized such that visually similar items are adjacent in a continuous visual feature space (e.g., a colour space). The advantage of this paradigm is that researchers can quantify the representational error that a cognitive system makes when processing the inputs by calculating the distance between the responded value and the actual target in the scale. Specifically, when a substantial number of errors are accumulated by repeating trials, they can typically observe a Gaussian shaped error distribution around the target, which is conceptually well-matched with internal noise assumed to arise during the representation process. That is, with an appropriate response scale, the error distribution can be interpreted as a reconstruction of our mental representation.

Since it provides direct reconstructions of visual representation, the continuous report paradigm has broadened our understanding of visual perception and memory. For example, researchers have quantified more detailed aspects of visual representations like memory precision or bias by quantifying how densely errors are distributed near the target or how the central tendency of a response distribution is systematically shifted (Oh et al., 2019; Son et al., 2020; Wilken & Ma, 2004). The VWM literature has leveraged a mixture model approach that teases apart different components of the error distribution that are theorized to reflect fundamental retention mechanism of VWM (Bays et al., 2009; Fougnie et al., 2010; van den Berg et al., 2012; Zhang & Luck, 2008; but see also Schurgin et al., 2020). Specifically, despite some variations, the basic structure of the mixture model consists of a von-Mises distribution that reflects the correct memory for the target with internal noise and a uniform distribution that accounts for pure guessing. By fitting this mixture model to the data, mechanistically relevant properties of error distributions can be identified, such as how often participants forget the stimuli or how precisely they can remember the target.

Despite the apparent advantages of the experimental paradigm based on highly controlled stimuli, to our knowledge no studies have attempted to use this approach to characterize perceptual and memory representations for realistic scene environments. The key challenge stems from the difficulty in stimulus control for real world scenes. To create a continuous stimulus space, it is necessary to parametrically manipulate the targeted properties of the stimuli (e.g., hue in colour wheel) such that changes occur without any abrupt transitions at a fine scale with constant increments. Since this level of control is very hard to achieve in real-world scenes, several studies have instead embraced continuous spaces of simple features in real-world stimuli (Brady et al., 2013; Miner et al., 2020; Chanales et al. 2020; Sun et al. 2017). For example, Sun et al. (2017) systematically manipulated colours of real-world objects and asked participants to remember and report the colour of the target object under a continuous report paradigm. Based on the mixture model, they are able to distinguish behaviour driven by complete forgetting of stimulus information (guessing) from less precisely remembered information (decrement in precision). They found that colour memory for a target object can experience both modes of failure depending on colour similarity between target and distractor objects. However, since colour is only one attribute of the object stimuli, findings from this approach are more likely to reflect colour memory itself rather than reflecting the integrative nature of object memory. Moreover, although this work represents a departure from typical VWM paradigms based on arrays of coloured squares or circles, it is unclear how continuous colour space manipulations can be extended to real-world scenes. Not only are real-world scenes composed of multiple objects, each with their own primary colour, but much of rapid scene perception and categorization has little to do with colour information and these processes can even be hampered when a scene’s major color scheme is abonormal (Castelhano et al., 2008; Delorme et al., 2000; Goffaux et al., 2004).

Conceptually, a continuous stimulus space for real world scenes should depend on the complex features critical for distinguishing between scene categories (Walther & Shen, 2014; Wilder et al., 2019). Characterizing such features has been a key question in scene perception for decades (Davenport & Potter, 2004; Greene & Oliva, 2009; Kauffmann et al., 2015; Oliva & Schyns, 2000; Oliva & Torralba, 2001; Schyns & Oliva, 1994; Walther et al., 2011); however, the recent revolution in deep learning approaches from machine learning and computer vision have provided novel insights into the nature of features underlying visual categorization (Cichy et al., 2017; Eberhardt et al., 2016; Krizhevsky et al., 2012; Rezanejad et al., 2019; Zhou et al., 2017; for review, Serre, 2019). In particular, generative adversarial networks (GAN; Brock et al., 2018; Goodfellow et al., 2014; Karras et al., 2019; Shocher et al., 2020; Yang et al., 2019) offer a data-driven method for uncovering the complex features spaces necessary for generating artificial but highly realistic images. GANs are composed of two deep neural networks, a generative network and a discriminative network, which perform antagonistic tasks. The generative network produces artificial images from random inputs while the discriminative network distinguishes real photographs from the artificial images synthesized by the generative network. These networks are placed in opposition with the generative network aiming to generate synthesized images that fool the discriminative network, and the discriminative network adapting to correctly categorize real from synthetic. Thus, the GAN is set to train the generative network so as to minimize the performance of the discriminative network. After successful learning, the synthesized images created by the generative network are highly realistic even to human observers. Recent studies of scene specific GANs (Shocher et al., 2020; Yang et al., 2019) have demonstrated that latent spaces from trained generative networks show a meaningful organization such that adjacent vectors in latent space are highly similar to each other and their associated synthetic images represent continuously changing realistic scenes. These advances offer a unique possibility – a continuous space of natural scene images may be constructed by deliberately traversing the latent space of a GAN trained to synthesize scene images.

In the current study, we test whether a scene space synthesized from a GAN can be used to generate continuous scene stimulus sets for the investigation of human perception and memory. To do this, we leveraged a GAN trained on natural scene images (Yang et al., 2019) and designed the way we select latent vectors such that the synthesized output results in a continuously changing scene space. Specifically, we sought to generate circular scene spaces, which we call scene wheels, akin to colour or orientation wheels that continuously traverse a stimulus space with no specified start and end points. To do this, we selected a set of latent vectors located on an imaginary circle in latent space and generated the associated synthetic scene images. In particular, we parametrically manipulated the radius of the circles in latent space to explore how the generated scenes vary depending on distances between their latent vectors, and how this variation affects human perception and memory performance. To assess and validate the continuous nature of changes throughout the scene wheels, we conducted a perceptual similarity judgment task with the scene images and compared it with pixel-wise correlations of the images. We anticipated that if our approach to generating scene wheels actually reflected the latent scene spaces fundamental to human scene perception, scene wheels with larger radii should be perceived as less similar. We found clear evidence of this pattern and also confirmed that this pattern is reflected in pixel-wise correlations of the scene images. In addition, we hypothesized that images on the larger wheel should be remembered with higher precision, because other images neighboring the to-be-remembered scene would interfere less with participants’ memories. To test this hypothesis, we conducted a VWM experiment using a delayed estimation task based on the continuous report paradigm. Participants successfully remembered and reconstructed scene images from the scene wheels in their WM, and the observed error pattens are consistent with typical error distributions observed from similar experiments with simple stimuli (Bays & Husain, 2008; Wilken & Ma, 2004; Zhang & Luck, 2008). Critically, the radius of the scene wheels was associated with memory precision. Our work provides a novel method to measure memory performance of realistic scene stimuli as well as a new class of highly controllable scene stimuli for a range of experiments on scene perception and memory.

## Methods

### Participants

#### Perceptual similarity judgment

One hundred and twenty eight participants (25 females, 101 males, 1 other, 1 missing, mean age = 34.88) were recruited online through Amazon Mechanical Turk. We excluded 28 of the participants from the main analysis based on the criterion explained in the *Analysis* section (screening rate = 21.87 %). This sample size was determined for the purpose of collecting a sufficient amount of data to validate all 25 wheels. All participants reported normal or corrected-to-normal vision and received monetary compensation for their participating.

#### Delayed estimation

20 participants (16 females, 4 males, 0 other, mean age = 23.31) with normal or corrected-to-normal vision were recruited from University of Toronto. The experiment was conducted online and all participants were compensated with monetary rewards. The targeted sample size was pre-determined based on a power analysis from a pilot experiment (N=17). The power analysis focused on the fixed effect of scene wheel radius as defined in a mixed effect regression model (see details in Analysis) and was conducted using the SIMR package (Green & MacLeod, 2015) in R 3.6.1 (R Core Team, 2019). A power curve was estimated by simulating 100 datasets and refitting the regression model to each simulated dataset for each of a range of sample sizes from 4 to 19 participants. The results showed that with a sample size of only 4, the estimated power reached 95% (CI [88.72, 98.36]). Thus, we decided to collect 20 participants for the main experiment, a similar number with the pilot experiment. The study protocol was approved by University of Toronto Research Ethics Board.

### Stimuli

The continuous scene space, or the scene wheel, was generated using the StyleGAN model trained with a mixed image pool of three indoor scene categories – bedrooms, living rooms, and dining rooms (Yang et al., 2019). The training image set included 500,000 scenes per scene category all of which were selected from the LSUN dataset (Yu et al., 2015). Generating scene wheels from this GAN required a systematic approach to sampling the latent vectors of the generative model. First, we randomly sampled three latent vectors and defined a plane by calculating the vector cross products of the three vectors. On that plane, an imaginary circle was drawn around the center point of the three seed vectors. We then selected 360 points equally spaced by 1-degree unit on the imaginary circle (Figure 1). The extracted points were fed into the generative network as input and synthesized into natural-looking scene images. Figure 2 shows the images synthesized by this procedure, including images representing 5 different center points and the images generated from one of those center points (see Supplementary video for all images at https://osf.io/h5wpk/).

**Figure 1.**
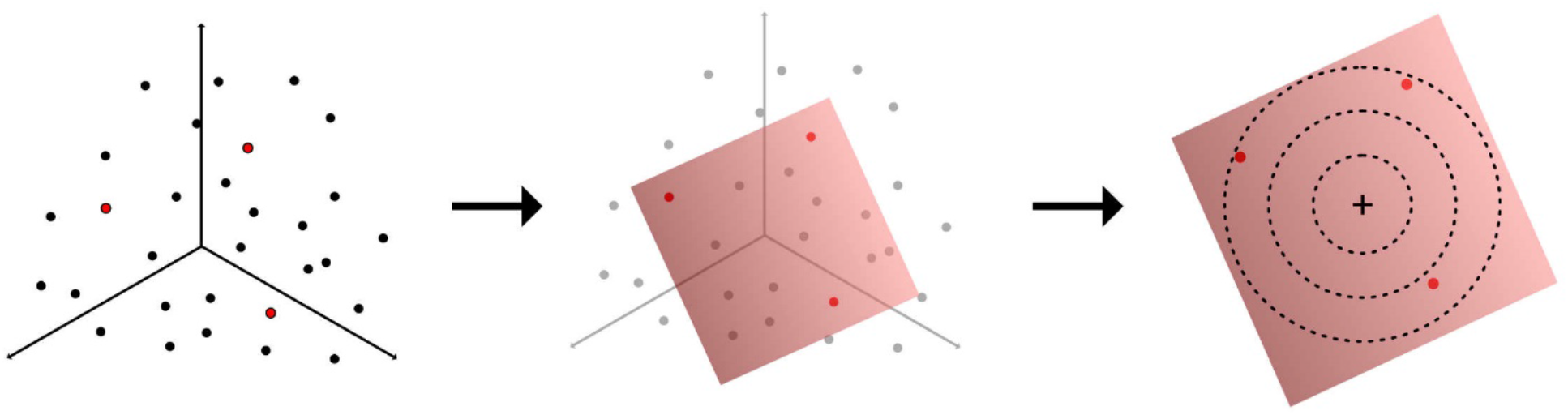
A simple scheme of wheel generation procedure. The selected points on the imaginary circles (third stage) were used as input to the generator.

We also examined how composite images of a scene wheel are affected by differences in radii of the wheel in the latent space. To do this, we created separate imaginary circles by repeatedly doubling the radius while maintaining the center of the circle. Scene wheels with smaller radii resulted in scenes with few changes in their visual properties (Figure 2B) whereas circles with larger radii contain less similar images with larger changes (Figure 2C). This pattern was highly consistent across scene wheels generated from different origins. We chose 5 different radii (2, 4, 8, 16, and 32 a.u.) to test whether perception and memory performance systematically changes with radius. Since we generated different types of wheels from 5 different origins, a total of 25 wheels were created.

**Figure 2.**
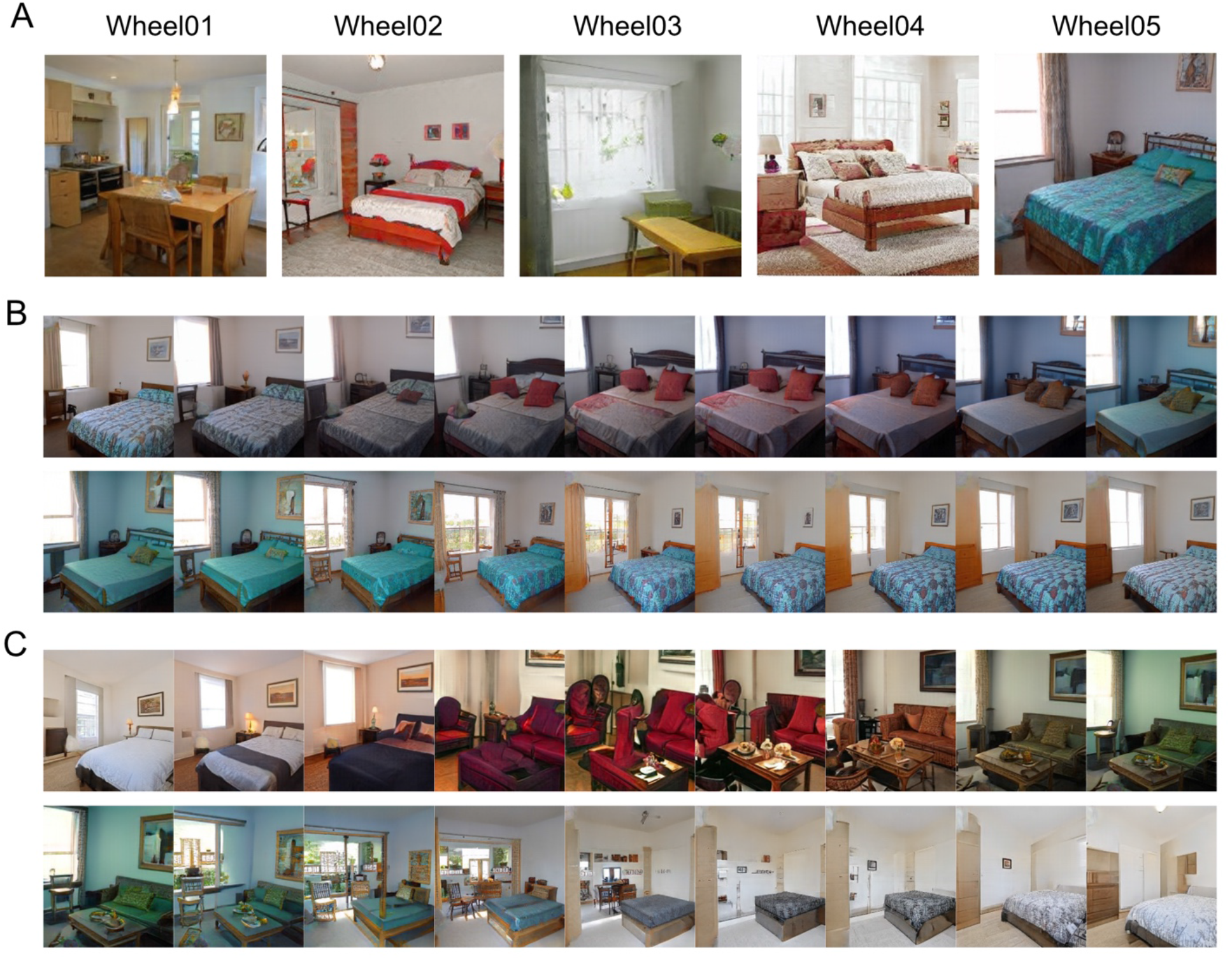
A) Images corresponding to the center points of the five wheels. B & C) Example images of two scene wheels that share the same centre point but vary in radius. The images in these panels are equally spaced by 20 degrees from 0 to 340 degrees. B) 18 example images from Wheel05 with radius of 8. The top-left and bottom-right scenes are located at neighboring locations on the wheel (0 and 350 degrees) thus are very similar. C) 18 examples from Wheel05 with radius 32. Compared to B, the changes between images are larger.

The wheels generated through this process were validated with behavioral experiments for perception and memory. For the perceptual validation, we sampled 12 scene images at 30-degree intervals and created 66 unique pairs in each wheel for the pair-wise similarity judgment task. We generated 360 images from each wheel in one-degree intervals for the working memory experiment using the continuous report paradigm. To test the full range of each scene wheel, we divided each wheel into 20-degree bins and randomly selected three scene images from each bin as targets for each participant. Due to the web-based nature of the study, the image stimuli were displayed at different sizes depending on the individual participant’s monitor settings. The images occupied 20% of the monitor width and were presented with a square aspect ratio. Note that the experiment was set to be run only on laptop or desktop computers, but not on smartphones or tablets.

### Procedure & Design

#### Perceptual similarity judgment

In each trial, a pair of scene images was shown side by side along with a 6-point Likert scale (Figure 3A). Participants were asked to click a score on the scale according to how similar they think the two images are (see the demo version of the experiment: https://macklab.github.io/sceneWheel_similarityJudgment/demo.html). Before starting the main session, participants completed a set of 10 practice trials using scene pairs from a wheel not used in the main session. Those scene pairs were selected to represent the full range of physical similarity (pixelwise correlation coefficients) so that participants could calibrate to the potential range of pair similarity. The full experimental design was 5 (between-subject variable: different center points) x 5 (within-subject variable: wheel radii) x 66 (image pairs per wheel, combination of 2 in 12 images with no replacement). 20 participants were recruited for each center point condition. Each participant completed 340 trials, which consisted of 330 main trials (66 image pairs x 5 radii condition) and 10 catch trials that showed identical images. The catch trials were used for screening participants who showed bad performance and were thus excluded from main analysis.

#### Delayed estimation

The procedure followed the conventional delayed estimation task in visual working memory (Wilken & Ma, 2004) with a small variation on display durations. Each trial began with a memory display with a target scene image (1 sec) and followed by a short retention interval for approximately 1 second. After the retention interval, a response display with a gray box and indicator wheel was presented. Once participants moved their computer mouse, the gray box was replaced with a random scene within the scene wheel. Thereafter, the scene shown changed according to the mouse position. By moving the mouse cursor along the indicator wheel, participants were asked to find a scene that closely matched the target scene presented in the memory display. Participants responded with their best matching scene candidate by clicking the left mouse button. To avoid effects of location memory, the probe scene wheel was randomly rotated in each trial. An example trial sequence is depicted in Figure 3B (demo experiment: https://macklab.github.io/sceneWheel_main/demo.html).

We used a mixed design with 5 (between-subject variable: 5 different center points) x 5 (within-subject variable: wheel radius) conditions. 4 participants were recruited for each scene wheel and each participant performed 270 trials (54 trials for all 5 radius levels of a single scene wheel). Before starting the main experiment, participants completed 5 practice trials. During the main experiment, short breaks were given every 54 trials.

**Figure 3.**
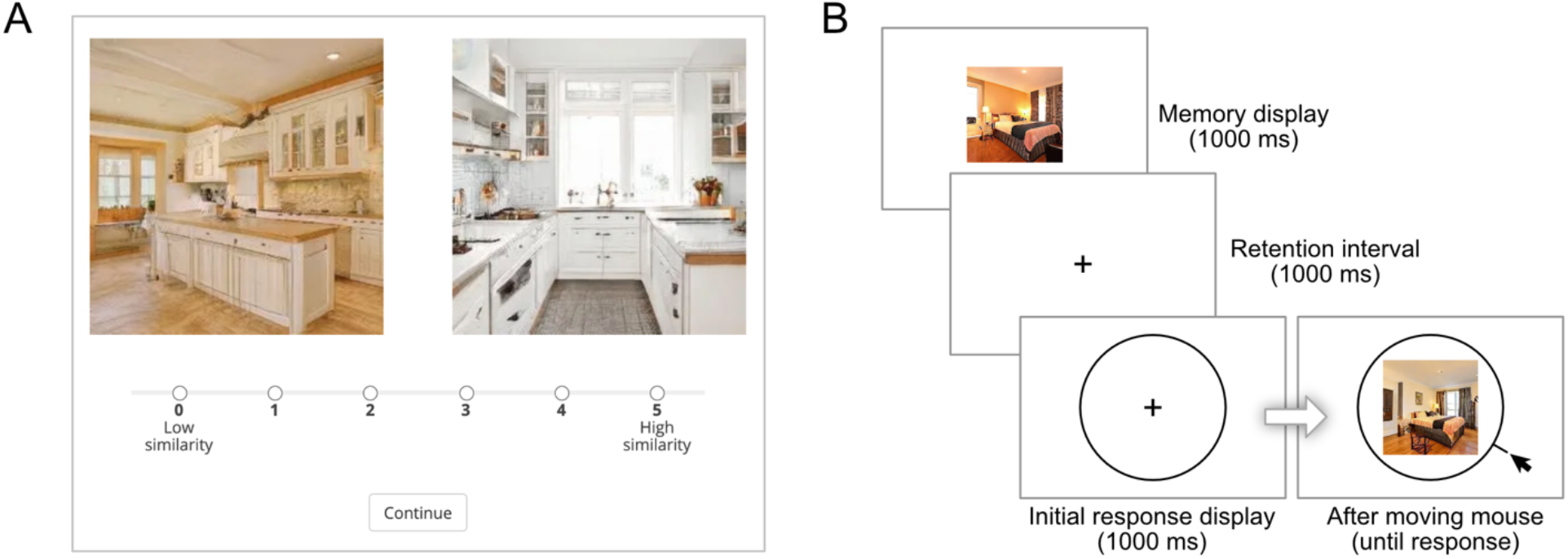
A. Example trial ofperceptual similarity judgment task. B. Example trial sequence of delayed estimation.

### Analysis

#### Perceptual similarity judgment

We quantified the perceptual similarity of the scene images on wheels to confirm the continuous nature of scene images along their position on the wheel. Before the analysis, we excluded 28 participants who showed poor performances on the task based on a two-step exclusion process. First, we analyzed the performance in the 10 catch trials, in which identical images were shown side by side. Participants who rated more than 70% of the catch trials below the score of 4 were excluded in this step. Second, we excluded data if the distribution of rating responses showed extremely large kurtosis over 5. These participants rated most of the trials with a single score. We excluded 28 participants in total, 14 in step 1 and 14 in step 2. The similarity scores of the final 100 participants were normalized to the range between 0 and 1 and averaged across the respective set of 20 participants who viewed the wheels with the same center point (Figure 4). For a direct illustration of the continuous similarity pattern, we sorted the mean similarity scores of all pairs according to their angular distance on the scene wheel (Figure 6A).

#### Pixel-wise correlation

For practical reasons, we obtained perceptual similarity ratings only for 12 images per scene wheel, spaced 30 degrees apart. Assessing physical similarity between the images, however, poses no such constraints. We quantified physical similarity between all 360 images in each wheel by computing their pixel-wise correlation (Figure 5). For better illustration, we again sorted the correlation coefficients of the pairs according to their angular distance on the scene wheel, by averaging all Fisher-z transformed correlation coefficients with the same pair distance (Figure 6B). We compared the physical and perceptual similarity scores by conducting Spearman’s rank correlation between the lower triangle of the perceptual similarity matrices and the corresponding values from the pixel-wise correlation matrices.

#### Delayed estimation

To examine whether memory performance corresponds to perceptual similarity changes among different radii, we measured memory response errors and calculated circular standard deviation (SD) of errors in each radius condition. First, response errors for each trial were calculated by subtracting the target angle from the angle of the response that participants made in each trial. After sorting the response errors into 5 radius conditions and 5 center point conditions, we calculated circular SDs from the response error distributions using the circular package (Agostinelli & Lund, 2017) in R 3.6.1 (R Core Team, 2019). Then, we applied a linear mixed-effects model that predicts circular SDs with a fixed effect of the wheel radii and the random effect of participants using the LME4 package (Bates et al., 2015).

## Results

### Perceived similarity pattern

To evaluate the perceptual similarity pattern of scene wheels, we collected similarity rating scores of all pairs of 12 composite scenes (66 pairs) in each wheel (Figure 4). Diagonal burgundy lines indicate image identity – these were not assessed in the experiment, except in 10 catch trials. Across all center point conditions, we consistently observed that images on the wheels with smaller radii were perceived as more similar, depicted by warmer colors. This result supports our anticipation that wheels with smaller radii contain more similar images because the corresponding latent vectors are located closer to each other. For each wheel, we also observed a graded color transition: cells closer to the diagonal line show higher similarity rating scores (warm colours), whereas farther cells show lower scores (cool colours). To further examine this pattern, we averaged the similarity scores of image pairs whose angular differences are the same on the wheel (Figure 6A). Across all center points with all radii, mean similarity rating scores of image pairs continuously decrease as their angular distances on the wheel increase, indicating that pairs of scenes become less similar the further apart they are located on the scene wheel. Moreover, the drop-off gets steeper for larger radii, again showing that similarity of scenes systematically varies with the wheel radius.

**Figure 4.**
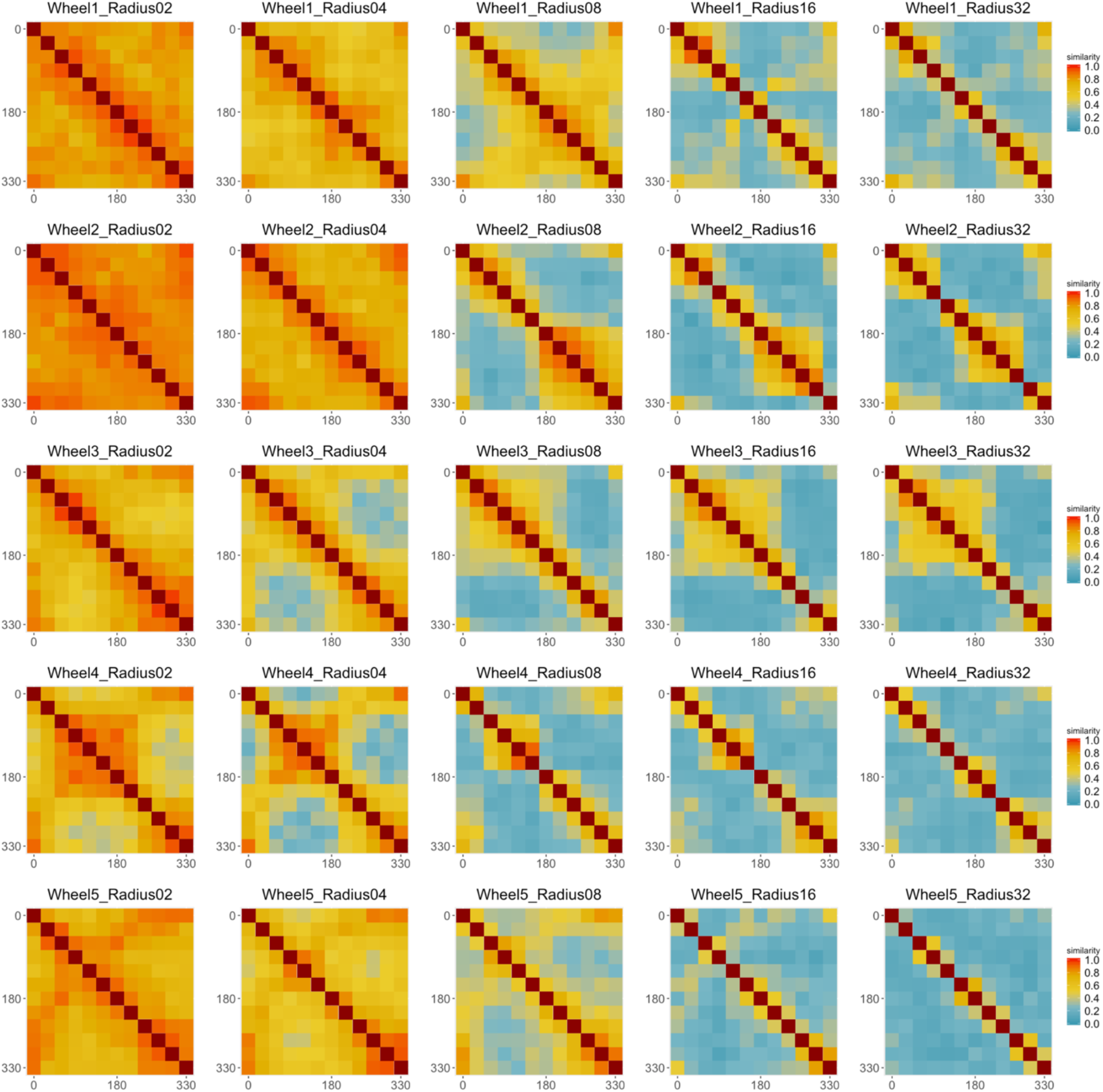
Perceptual similarity matrices. Each row indicates wheels with a different center point. Each column corresponds to a different radius. Across center points, smaller radii show higher correlation depicted by warmer color.

### Pixel-wise correlation

Figure 5 shows the pixel-wise correlation matrices of all scene wheels. Similar to the perceptual similarity matrices, we again observed that images on the wheels with smaller radii and closers to the diagonal show higher correlation coefficients. Figure 6B shows the mean coefficient values sorted by the angular differences. We could observe the gradual decrease of correlation coefficients, comparable to the pattern of perceived similarity scores in the upper panel. This comparability between the physical and perceptual similarity of the scene wheels was confirmed by a high correlation between the lower triangles of the similarity matrices of the two measurements, averaged across all wheels (mean Spearman’s *ρ* = 0.63; mean computed under Fisher’s-z transformation).

**Figure 5.**
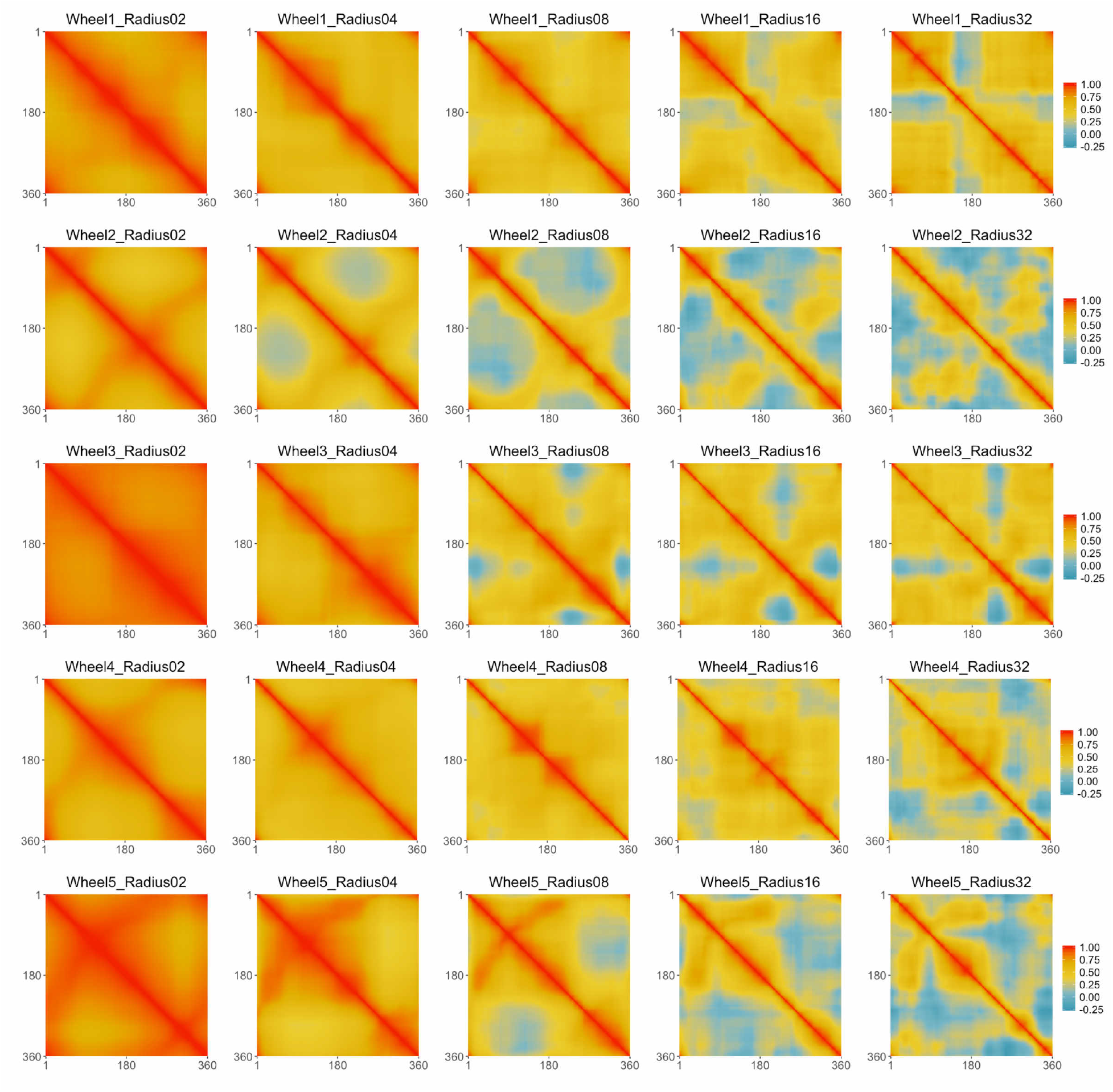
Pixel-wise correlation matrices. Each row shows the correlation results of wheels with different center points. Each column corresponds to a different radius. Across center points, smaller radii show higher correlation depicted by warmer color.

**Figure 6.**
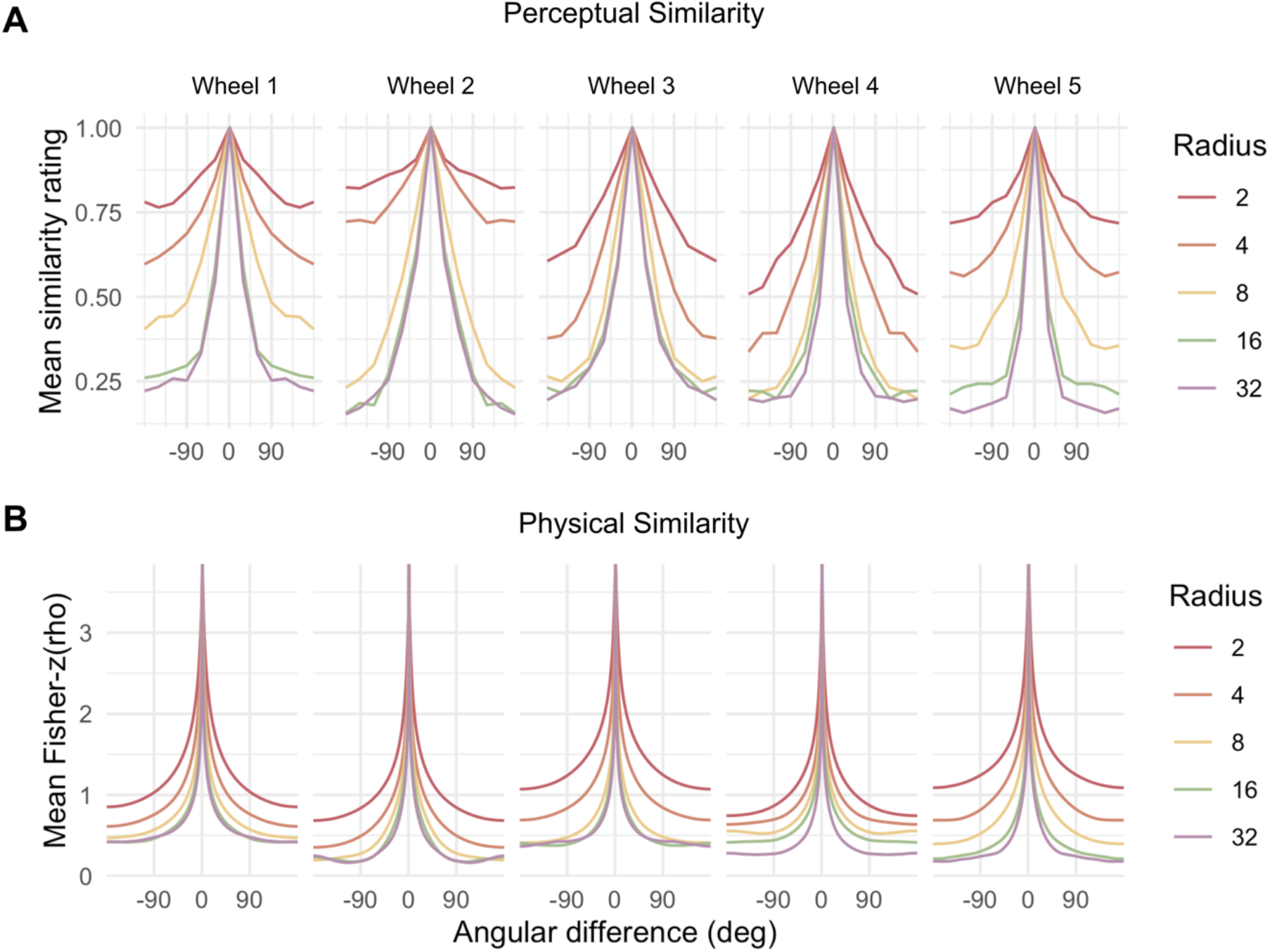
A. Mean perceptual similarity rating scores aligned by the angular differences on each scene wheel. Values at 0 degrees (identical pairs) are set to one. B. Mean correlation coefficients aligned by the angular difference within each scene wheel. In both A and B, similarity values drop off as the angular differences get larger across all wheels. The steepness of the drop-off depends on the wheel radius.

### Working memory performance

We tested if the changes in physical and perceptual properties of scene wheels confirmed in the previous section are reflected in human memory. Specifically, we conducted the VWM experiment using the continuous report paradigm to test if scene wheels with larger radii where composite scenes look less similar are remembered more precisely. The error distributions of each radius condition are provided in Figure 7A. The observed error distributions resemble Gaussian distributions, as reported in previous studies using the continuous report paradigm and simple stimuli (Husain & Bays, 2009; Wilken & Ma, 2004; Zhang & Luck, 2008; see *Direct comparison to previous studies* section for details). We then applied a linear mixed-effects model to the data to predict memory precision represented by circular SDs of errors with radii of the wheels, since we hypothesized that memory precision would increase with larger radius. As expected, we observed that a strong effect of the radius on circular SDs (*F*(1, 79) = 205.32, *p* < .001), suggesting that memory precision increases with the increment of wheel radius (note that smaller circular SD indicates higher memory precision). Then, we further explored the effect of the center points to see if it affects the memory precision in addition to the radius by estimating a linear mixed model that included the interaction between the wheel type and radius. Relative to the first model with only the radius factor, we found that the interaction model explained the data better (χ^2^(8) = 21.48, *p* = .006). The interaction of the wheel type and radius was significant (*F*(4, 75) = 4.22, *p* = .004), which is reflected in slightly different slopes of the wheel types (Figure 6B). Despite these slight differences in the magnitude of the effect across wheel types, we found that participants’ memory for the realistic scenes was more precise with larger radii, and this increasing pattern of memory precision along the increment of radius is not limited to specific wheels but a common effect that can be applied to all wheels.

**Figure 7.**
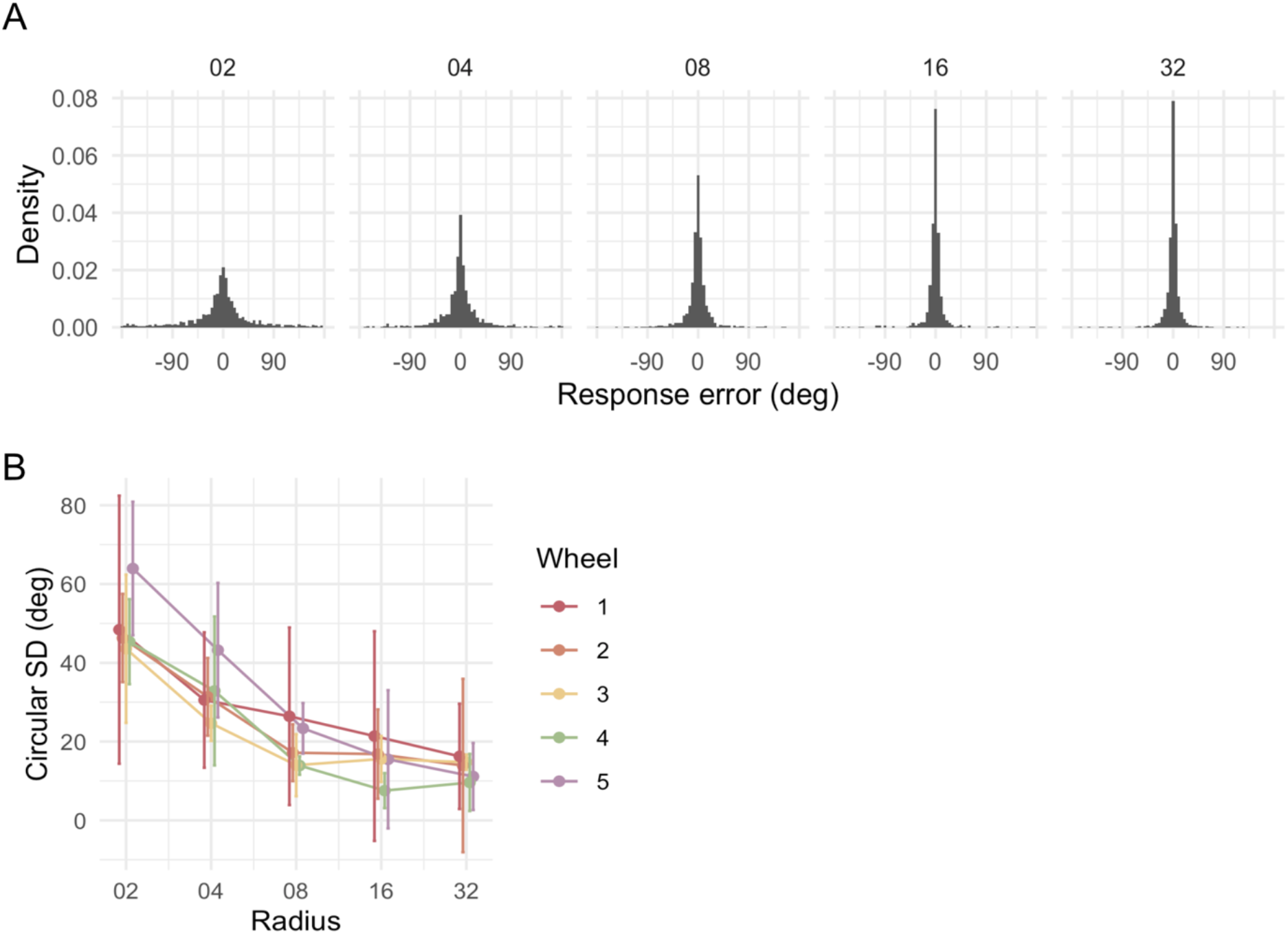
A. Histograms of response errors in each radius condition. All participants’ data and corresponding wheel type conditions are aggregated. Density of the histograms decreased with increasing radii (from left to right panel). Bin width is 5 degrees. Individual participant data are available in the Supplementary Figures. B. Decreasing pattern of circular SDs of the response errors with increasing radius, indicating that memory precision increases with increasing radius. Error bars: 95% confidence intervals.

### Direct comparison to previous studies

We further examined the resemblance of the changes in circular SDs between our study and previous studies conducted with simple stimulus domains (Bays & Husain, 2008; Wilken & Ma, 2004; Zhang & Luck, 2008). These studies reported gradual changes in the circular SDs of error distributions with increasing set size (the number of items held in VWM), a factor commonly utilized to control the difficulty of memory experiments. In our study, we also found a gradual change of circular SDs with increasing radii of the scene wheels, which served as a similar control of task difficulty. Thus, we checked how the circular SDs of our error distributions relate to the circular SDs observed in the previous studies. We found that the range of circular SDs in our study spans the range of the previously reported SDs (Figure 8). We conclude that the error distributions obtained using scene wheels resemble the error distributions from other, simpler stimulus domains.

**Figure 8.**
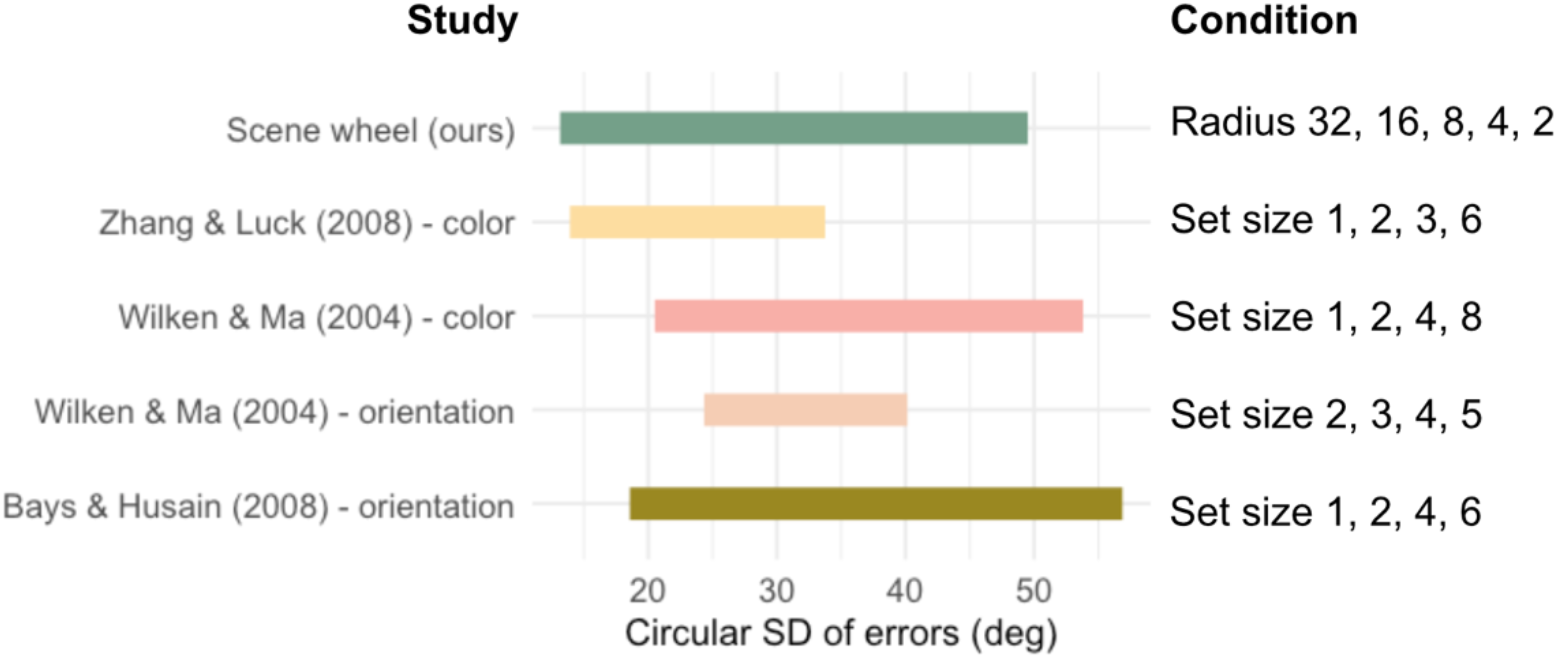
Illustration of observed error SD ranges. The range of the current study (top row, scene wheel) spans the ranges of error SDs observed by the other studies.

## Discussion

We demonstrated that continuous scene spaces generated with a GAN provide a novel tool to measure visual perception and memory performance for complex realistic scenes. Specifically, we confirmed that by carefully selecting latent vectors from a pretrained GAN, the similarity among scene images can be well controlled, allowing for a continuously changing traversal of meaningful scene space. With the generated scene wheels, we validated the perceived similarity of the generated scene images using pairwise similarity judgment. We observed that participants’ similarity judgments corresponded to the images’ distance on a wheel. Also, this perceived similarity pattern was highly correlated with pixel-wise correlation, suggesting that pixel-wise correlation provides a proxy to examine the continuous nature of newly generated wheels according to perceptual similarity without further empirical validation. For mnemonic validation, we conducted a visual working memory experiment using the continuous report paradigm. We observed that the error distribution from the participant’s responses was comparable with the typical error distribution that previous studies observed from simple stimuli like colours. Critically, we additionally manipulated the radii of the wheels in the latent space to parametrically control the similarity among the scenes in the wheels. We found clear evidence that participants’ memory systematically varied with this manipulation of scene similarity as predicted. Therefore, we suggest that this novel method to generate scene stimuli using a GAN not only allows for an unprecedented level of stimulus control for complex scene stimuli, but also that the latent spaces generating our scene wheels reflect fundamental representational spaces important for human scene perception and memory.

As is evident in the perceptual similarity measures, pixelwise correlation, and memory validation study, the latent GAN space is complex with regions of inhomogeneity. This means that scene wheels generated from different center points will likely include distinctive scene properties that uniquely impact perceptual and mnemonic similarity. It is important to consider this limitation if the empirical goal is targeting perfect generalization of specific effects across different scene wheels. However, despite these small scene wheel-specific variations, we observed a reliable and consistent pattern of perceptual and mnemonic performance on different scene wheels. We anticipate that the level of control achieved with the stimuli generated from the GANs will be valid for the majority of experiments investigating perception and memory of real-world scenes.

Based on this level of control over the scene stimulus space, we expect that findings from using simple stimuli, such as colour wheels, can be generalized to photo-realistic scenes. First and foremost, our approach can be leveraged to measure memory performance for scenes at a fine level of granularity that was not available previously for realistic scenes. In the recognition paradigm, for example, our methods could be utilized to parametrically control the perceptual similarity between target and foil scenes to systematically test how scene similarity affects memory interference or confusion (Konkle et al., 2010; Lin & Luck, 2009). In using a continuous report paradigm with the GAN-based scene wheels, as we did here, one can measure specific components of scene memory, such as precision or bias, on a continuous scale. Coupling the current approach with cognitive models like the mixture model allows for quantitative estimates of memory capacity and errors for scenes in the same way conventionally done for simpler stimuli (Bays et al., 2009; Brady & Alvarez, 2011; Fougnie et al., 2010; Son et al., 2020; Sun et al., 2017; van den Berg et al., 2012; Wilken & Ma, 2004; Zhang & Luck, 2008). Additionally, the continuous scene space can be utilized in any paradigm that requires carefully controlled stimuli. Given that simple stimuli and realistic scenes have notable differences in their complexity and the amount of information, comparing the results from those levels of stimuli will contribute to elaborating how memory processes change along the visual hierarchy and provide testable predictions for neural function and representation.

The scene space from the GAN has the potential to achieve even more deliberate control over particular scene properties. Recent studies showed that different layers of the generative network in GANs represent different properties of the scenes (Karras et al., 2019; Schocher et al., 2020; Yang et al., 2019). By manipulating the latent vectors in a layer-wise manner, one can exclusively control specific properties of the scenes that the targeted layer addresses, while the other properties remain the same. For example, Yang et al. (2019) suggested that a visual hierarchy emerges in layers of the generative network such that low-level scene properties like layout are processed in lower layers while higher-level properties like detailed attributes of the scene (e.g. composite objects, materials of the objects, lighting condition of the scenes, etc.) are processed in higher layers. By manipulating the latent vector in certain layers, GANs are able to synthesize a set of scenes with parametrically different attributes (e.g. lighting, amount of clutter) but with limited changes in the other properties (e.g. layout and category). Based on this layer-wise manipulation, future studies may investigate various level of information embedded in realistic scenes, while minimizing effects from uncontrolled properties.

Our study is not the first attempt to generate realistic stimuli on a continuous scale. Previous studies took an approach to morph stimuli with assigning different weights to each reference stimulus in the various stimulus domains such as animal species (Freedman et al., 2001; 2003), real-life objects (Newell & Bülthoff, 2002), cars (Folstein et al., 2012; Jiang et al., 2007), or faces (Beale & Keil, 1995; Goldstone & Steyvers, 2001; Haberman & Whitney, 2007; Steyvers, 1999). This approach allowed participants to respond on a continuous scale similar to our study. However, to get a continuous interpolation between morphing references, the reference images need to be similar and well matched in terms of their physical status. For example, to continuously manipulate the emotion of faces, reference faces should be closely matched in properties like identity, gender, and spatial configuration of the facial components (e.g. eyes, nose, and lips), which requires time-consuming preprocessing by researchers to utilize the morphing algorithm per se. For this reason, the morphing approach was restricted to stimulus domains with little physical variance among the exemplars (e.g. faces have fewer physical variances than scenes). Our approach using a GAN, however, can be applied to highly complex naturalistic scenes since it utilizes neural networks trained on a large set of photographs and thus creates a large latent space of possible images. That is, each GAN’s latent space itself can be considered as a huge geometrically structured image database with high controllability. Synthesizing realistic images using the GANs, therefore, will expand the range of stimulus domains that can be successfully adapted in visual cognition studies.

Some recent studies in the visual memory field have adapted a different measurement paradigm, free recall, to target memory precision (Bainbridge et al., 2019; Bainbridge & Baker, 2019). In this paradigm, as conveyed by its name, participants are asked to freely recall whatever they can remember after seeing the scene images by drawing the scenes. Since this paradigm does not require foil images to be compared to the target representation in memory, it measures memory representations more directly and provides insights into the details of the nature of scene memory, such as what information is remembered from the scene, and how much detail survived during retention. Nevertheless, this paradigm has practical challenges. Drawing is a qualitative measurement that needs to be scored using intrinsically subjective judgments for use as a quantitative measurement. This means that the measurement procedure in this paradigm requires at least two steps, drawing and assessment of drawing by a third person. Considering that the assessment is usually conducted by at least hundreds of participants, this is a labor-intensive paradigm. The other issue is that the drawing behavior itself could confound accurate measurement of memory performance. For many people, drawing is cognitive demanding requiring cognitive resources other than visual memory such as active hand movement or visual attention to the elements currently drawn, which could affect the retrieval process. Also, since drawing a complex scene often requires a substantial amount of time, the memory representation could suffer from memory decay. We believe that the technique proposed here can be another option for a fine-grained measurement of scene representation with more practical benefits.

Scenes generated from the GANs, however, can have a drawback in terms of synthesis quality. Depending on the pretrained GAN’s performance, synthetic scenes can sometimes include artifacts or “nonsense” regions (although in our experience, such scenes remain highly realistic and interpretable). Thus, it is key to first select well-trained GAN models and manually evaluate the generated scenes in preparation for conducting an experiment. We provide our scene wheel sets that are already tested with human participants in the current experiment (https://osf.io/h5wpk/) to help researchers to skip this procedure.

In conclusion, we anticipate that the continuous scene space approach proposed here will quickly be adopted as a fundamental tool for measuring scene representations. Going beyond the working memory experiment demonstrated here, we foresee that the scene wheels can be utilized in any domain that needs fine measurement and adjustment methods for realistic scenes. In leveraging this approach, we anticipate that future work will provide key insights into how we perceive and remember the real-world naturalistic environments that serves as the backdrop to our everyday experiences.

## Author Note

This research was supported by Natural Sciences and Engineering Research Council (NSERC) Discovery Grants (RGPIN-2017-06753 to MLM and RGPIN-2020-04097 to DBW) and Canada Foundation for Innovation and Ontario Research Fund (36601 to MLM).

The datasets generated during and/or analyzed during the current study are available in the Open Science Framework repository (https://osf.io/h5wpk/) and none of the experiments was preregistered.

The authors declare no conflict of interest.

## Supplementary Figures

**FS1.**
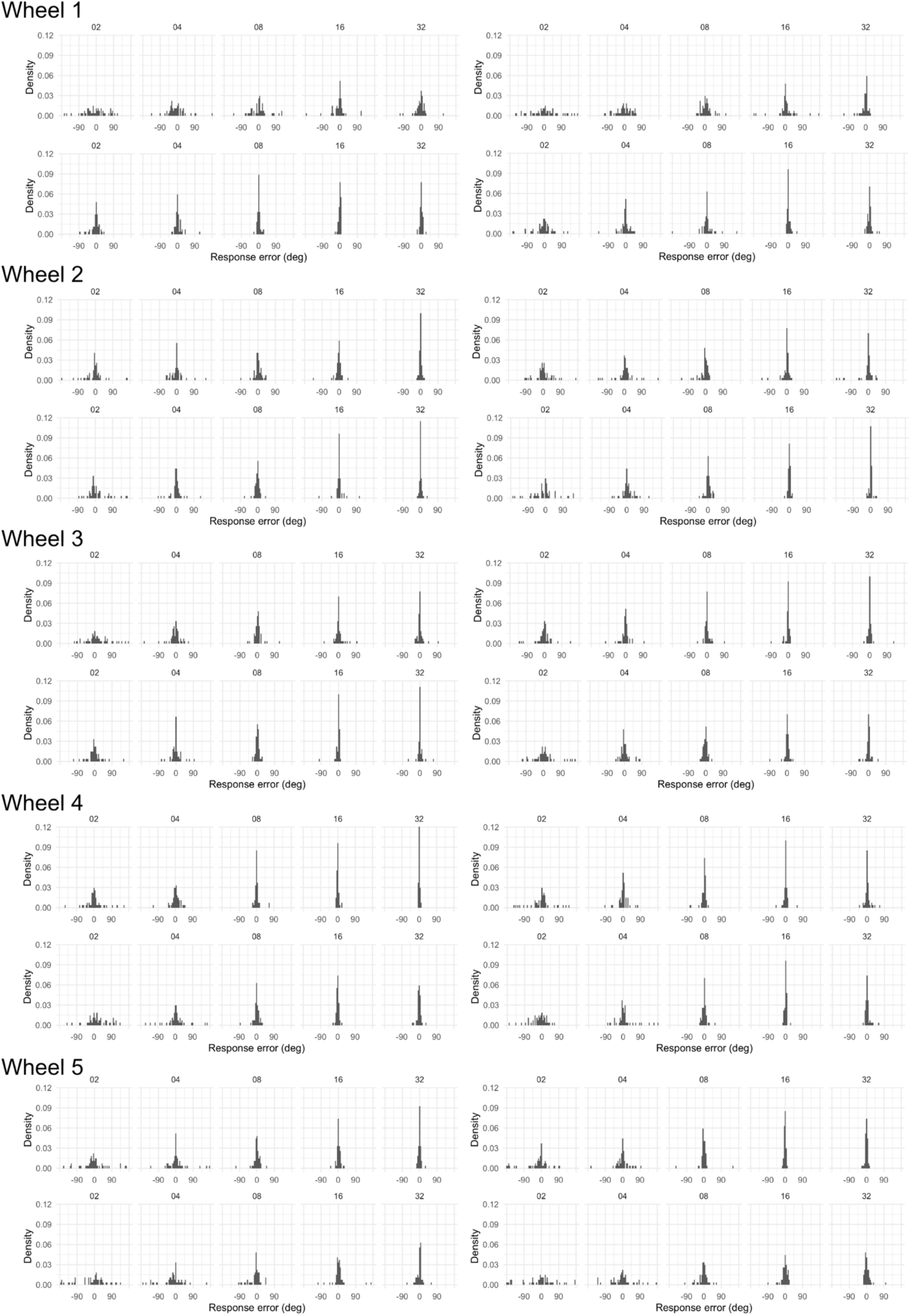
Error distributions of individual participant. Each panel containing five histograms indicates individual participant data. Across all wheel types and all participants, error distributions are highly similar, suggesting very consistent effects of radius condition. Bin width is 5 degrees.

